# MERS-CoV recombination: implications about the reservoir and potential for adaptation

**DOI:** 10.1101/020834

**Authors:** Gytis Dudas, Andrew Rambaut

## Abstract

Recombination is a process that unlinks neighbouring loci allowing for independent evolutionary trajectories within genomes of many organisms. If not properly accounted for, recombination can compromise many evolutionary analyses. In addition, when dealing with organisms that are not obligately sexually reproducing, recombination gives insight into the rate at which distinct genetic lineages come into contact. Since June, 2012, Middle East respiratory syndrome coronavirus (MERS-CoV) has caused 1106 laboratory-confirmed infections, with 421 MERS-CoV associated deaths as of April 16, 2015. Although bats are considered as the likely ultimate source of zoonotic betacoronaviruses, dromedary camels have been consistently implicated as the source of current human infections in the Middle East. In this paper we use phylogenetic methods and simulations to show that MERS-CoV genome has likely undergone numerous recombinations recently. Recombination in MERS-CoV implies frequent co-infection with distinct lineages of MERS-CoV, probably in camels given the current understanding of MERS-CoV epidemiology.

## Introduction

Recombination is an important process which expedites selection in many organisms (Muller, 1932) by unlinking loci. It also leads to different parts of recombining genomes to have different histories which, if not properly accounted for, can interfere with many genetic analyses, of which phylogenetic methods are amongst the most sensitive. Not accounting for recombination in phylogenetic analyses leads to incorrect (Schierup and Hein, 2000) and poorly supported genealogies (Posada and Crandall, 2002) and false inference of selection (Anisimova et al., 2003; Shriner et al., 2003).

With rising sequence availability during outbreaks of viral infectious disease, phylogenetic methods have been used to supplement our knowledge of epidemics in real time (Smith et al., 2009; Rambaut and Holmes, 2009; Lemey et al., 2009; Drosten et al., 2013; Cotten et al., 2013, 2014; Drosten et al., 2014; Gire et al., 2014). For some outbreaks there is little reason to suspect recombination, *e.g.* negative sense single stranded RNA viruses are thought to recombine over evolutionary, not population-level, time scales (Chare et al., 2003). Observable recombination in RNA viruses requires that two conditions are met: that viruses from distinct lineages co-infect a host and that a mechanism for recombination exists. For example, even though influenza A virus co-infection is extremely common in birds based on genome segment reassortment patterns (Li et al., 2004; Dong et al., 2011; Lu et al., 2014), recombination is extremely rare or absent (Chare et al., 2003; Boni et al., 2010). This is thought to be because template switching (Kirkegaard and Baltimore, 1986; Baric et al., 1987), the main mechanism of recombination in RNA viruses, is mechanistically difficult for single stranded negative sense RNA viruses (see Chare et al. 2003), and for influenza A viruses has only been convincingly shown in cell culture under extreme conditions (Mitnaul et al., 2000). When the genomic architecture of a virus is permissive to recombination, *i.e.* template switching occurs and is detectable, the extent of recombination is informative of co-infection and/or duration of infection.

Here we focus our attention on the Middle East respiratory syndrome coronavirus (MERS-CoV) (Zaki et al., 2012), a recent zoonotic infection with a relatively high case fatality ratio (Cauchemez et al., 2014; Memish et al., 2013; Assiri et al., 2013). Most human infections with MERS-CoV are thought to be the result of contact with *Camelus dromedarius* L., the dromedary camel, which is the presumed host of the virus. MERS-CoV, much like Severe acute respiratory syndrome coronavirus (SARS-CoV), is likely ultimately derived from bats (Corman et al., 2014a). MERS-CoV, along with Murine hepatitis virus and SARS-CoV, belongs to the Betacoronavirus genus. Betacoronavirus, as well as two other genera (Alpha- and Gammacoronavirus) out of four within the subfamily *Coronavirinae* have been shown to recombine in cell culture, *in vivo* and in eggs (Lai et al., 1985; Makino et al., 1986; Keck et al., 1988; Kottier et al., 1995; Herrewegh et al., 1998). Additionally, a coronavirus lineage related to MERS-CoV which was isolated from bats appears to have recombined around the spike (S) protein (Corman et al., 2014a). In this paper we show that although the genome of MERS-CoV contains considerable amounts of rate heterogeneity between genomic regions that can interfere with detection of recombination, we do nonetheless find evidence of sustained recombination that cannot be explained by rate heterogeneity alone. This has two important consequences: one is that care has to be taken when constructing phylogenetic trees of MERS-CoV as a single tree cannot accurately describe the complete history of all loci within a recombining genome. Secondly and more importantly, the observed rates of recombination in the MERS-CoV genome is evidence of a large number of MERS-CoV co-infections in some hosts which has implications for understanding the dynamics of the virus in the animal reservoir.

## Methods

### Overview

Recombination leaves several characteristic clues in genomes:

- Alternative topologies (Robertson et al., 1995a,b; Holmes et al., 1999). In some scenarios, for example if there has been a single large-scale recombination event, it is possible to clearly identify recombining fragments based on phylogenetic incongruity. Recombination can be inferred by reconstructing two or more phylogenetic trees from a partitioned alignment and looking for topological incongruity between them. Strong support for at least 2 incompatible phylogenetic trees across well-defined breakpoints is usually the most convincing evidence of recombination.
- Excessive homoplasies (Maynard Smith and Smith, 1998). The transfer of genetic material from one genetic background to another will result in apparent repeat mutations in different parts of a phylogenetic tree. However, it is possible for the same locus to undergo mutation independently, especially if the locus in question is under Darwinian selection. Detecting homoplasies alone is not sufficient to infer recombination, but should be demonstrated to occur in excess of expectation.
- Linkage disequilibrium (LD) decay (Meunier and Eyre-Walker, 2001). Linkage disequilibrium or LD is the non-random association of alleles at different loci. This is a statistic often reported for contemporaneous sequence data. In clonally (*i.e.* non-recombining) evolving organisms every allele is linked to every other allele in the genome and requires mutation to break linkage. In recombining organisms there is an expectation that LD will decay with distance between the loci, *i.e.* that loci further away from each other are more likely to be unlinked via recombination.

We test for each of these hallmarks of recombination in the MERS coronavirus genome using a combination of phylogenetic and linkage disequilibrium metrics. For a more detailed review of recombination detection methods see Posada et al. (2002).

### Alternative topologies

We use the Genetic Algorithm for Recombination Detection (GARD) method (Kosakovsky Pond et al., 2006), as implemented in the software package HyPhy (Pond et al., 2005), to look for alternative tree topologies in sequence data. Briefly, the method compares a model where a single tree is derived from the whole alignment and alternative models where breakpoints are introduced into the alignment and phylogenetic trees are derived independently from the resulting fragments. The presence of recombination, especially if it is recent and concentrated in some parts of the alignment, will result in two or more phylogenetic trees fitting the data better than a single tree model. It is important to note that likelihood estimation also involves other parameters, such as branch lengths, not just topology. We use GARD under a GTR (Tavaré, 1986) substitution model with Γ_4_-distributed rate heterogeneity amongst sites (Yang, 1994) on a dataset of 85 MERS-CoV sequences. GARD was run repeatedly until no more breakpoints could be identified in the resulting fragments.

In addition to this test, we run BEAST (Drummond et al., 2012) on partitioned coding sequences derived from the first well-supported breakpoint inferred by GARD. We extracted the coding sequences from nucleotide positions 1-23722 and 23723-30126 (referred to as fragment 1 and 2, respectively) of MERS-CoV genomes. Independent HKY+Γ_4_ (Hasegawa et al., 1985; Yang, 1994) nucleotide substitution models were specified for codon positions 1+2 and 3 and the analyses were run under an uncorrelated relaxed lognormal clock with an uninformative CTMC reference prior (Ferreira and Suchard, 2008) on the substitution rate for 100 million states, subsampling every 10000 states. The molecular clocks and trees of each genomic partition were either linked or unlinked, giving a total of 4 models. We used the multi-locus skygrid (Gill et al., 2013) as the demographic model for all analyses. Path-sampling and stepping stone sampling (Baele et al., 2012) were used to calculate marginal likelihoods and test the fit of each of the 4 models, under default parameters. In addition, 4 similar analyses were set up, but with strict molecular clocks, in order to contrast the performance of relaxed molecular clocks.

### Excessive homoplasies

Testing for recombination by looking for homoplasic mutations in phylogenetic trees requires that two conditions are met. One, that recombination is rare enough, so that there is sufficient phylogenetic signal to reconstruct the “correct” phylogeny otherwise known as the clonal frame (Milkman and Bridges, 1990). Two, that alternative explanations for homoplasic mutations can be dismissed with some certainty. There is no straightforward way of testing for the former, but the latter is usually dictated by the underlying biology. For example, repeat amino acid substitutions are a well documented response of influenza viruses and HIV to drug treatment (Gubareva et al., 2001; Tisdale et al., 1993; Boucher et al., 1993).

We employ two methods to test for excessive homoplasies. First, we use a maximum likelihood phylogeny inferred using PhyML (Guindon and Gascuel, 2003) under a GTR+Γ_4_ (Tavaré, 1986; Yang, 1994) nucleotide substitution model to recover a single tree using a MERS-CoV dataset comprised of 85 sequences. We then reconstruct ancestral sequences at each internal node and identify the mutations that have taken place along each branch using ClonalFrameML (Didelot and Falush, 2007). Mutations are then classified as either synapomorphies, shared variation derived via common descent or apparent homoplasies, shared variation derived from convergence, depending on how many times a given mutation has arisen in the phylogeny. The drawback of this method is that it necessarily conditions on a single tree with the highest likelihood.

We also employ BEAST (Drummond et al., 2012) to circumvent the limitation of conditioning the ancestral state reconstruction on a single tree. In addition to sampling various phylogenetic parameters from the posterior distribution BEAST is also able to map substitutions onto the branches of each MCMC-sampled phylogeny (O’Brien et al., 2009). This method is thus capable of estimating the posterior probability of a given mutation being synapomorphic or homoplasic by integrating over different tree topologies. Homoplasy analyses were performed on the concatenated coding sequences of MERS-CoV after partitioning the alignment into all 3 codon positions, each with an HKY nucleotide substitution model (Hasegawa et al., 1985) and no Γ-distributed rate heterogeneity amongst sites. A relaxed uncorrelated molecular clock with lognormally distributed rates (Drummond et al., 2006) under a CTMC reference prior (Ferreira and Suchard, 2008) and the flexible multi-locus skygrid as the demographic model (Gill et al., 2013) were used. The MCMC chain was run for 100 million steps, sampling every 10000 steps.

Throughout the paper we will refer to the number of branches that have experienced a given mutation as homoplasy degree. We define the homoplasy degree to be the number of times a given mutation has originated independently minus one. For example a homoplasy degree of 1 indicates that a mutation has occured on 2 different branches in the phylogeny. That is, we assume that one of the mutations has arisen through replication error, whereas the other has potential to have been introduced via recombination and thus can be thought of as excessive. Synapomorphies, on the other hand, are states that are shared by two or more taxa through common descent and thus necessarily are those mutations that have occured exactly once in the phylogeny. They have a homoplasy degree of 0 in all figures.

Additional tests for recombination were also performed, namely the estimation of the pairwise homoplasy index (PHI) (Bruen et al., 2006) and the triplet test implemented in 3Seq (Boni et al., 2007).

### Linkage disequilibrium decay

In the absence of recombination every allele should exhibit a high degree of linkage with other alleles in the genome. Under two extremes - clonal reproduction without recombination and free recombination - there is no correlation between LD and genomic distance and loci should be interchangeable. This is the basis of several non-parametric permutation tests for recombination that are implemented in the software package LDhat (McVean et al., 2002), which we used in combination with a dataset of 109 MERS-CoV genomes. Other, more complicated tests, such as composite likelihood methods, are also available but in our experience were incompatible with temporal sampling and rate heterogeneity.

### Sequence simulations

To test the performance of some of the methods we simulated two sets of sequences. We use fastsimcoal2 (Excoffier et al., 2013) to simulate 10 replicate datasets that have the same dates of isolation and similar diversity to the MERS dataset with 85 sequences under no recombination.

Additionally, we use πBUSS (Bielejec et al., 2014) to simulate sequences down an MCMC-sampled phylogeny drawn at random from a linked-tree unlinked-clocks BEAST analysis described above. We modelled region-specific rate heterogeneity by simulating a 30kb “genome” and setting the molecular clock rate for the first 20kb to be 9.5 × 10^−4^ substitutions site^−1^year^−1^ and the last 10kb to be 2.85×10^−3^, 1.9×10^−3^ or 1.3×10^−3^ substitutions site^−1^year^−1^, corresponding to roughly 3-, 2- or 1.3-fold rate heterogeneity between the two parts of the simulated genome. Two replicate datasets were generated for each category of rate heterogeneity. Other than that all simulations were run under a relaxed lognormal molecular clock (Drummond et al., 2006) with standard deviation set to 7.42×10^−7^, HKY substitution model (Hasegawa et al., 1985) with the transition/transversion ratio parameter (*κ*) set to 6.0 and Γ-distributed rate heterogeneity with 4 categories and shape parameter 0.04 and empirical nucleotide frequencies, all derived from the results of the marginal likelihood analyses described earlier. A MERS-CoV sequence isolated from a camel (NRCE-HKU270) was provided as the starting state at the root. To include the effects of site-specific constraint we additionally carried out simulations under a Goldman-Yang codon model (Goldman and Yang, 1994) in πBUSS with empirical codon frequencies, *κ*=6.0 and dN/dS (*ω*) set to 0.1 (*i.e.* purifying selection) under same levels of rate heterogeneity and on the same phylogeny as the simulations described above. As these sequences were simulated on a tree of MERS-CoV, we refer to these datasets as being empirically simulated. We also reconstructed ancestral states for these sequences using ClonalFrameML, as described above, to arrive at a null expectation for the number of homoplasies we expect to observe under rate heterogeneity but without recombination.

### Investigating the effects of temporal sampling and rate heterogeneity

All 10 sequence datasets simulated with fastsimcoal2 and 12 sequences empirically simulated in πBUSS were analyzed using LDhat (McVean et al., 2002) to ascertain the effects of temporal sampling, and in the case of πBUSS-simulated sequences, the effects of rate heterogeneity in the presence of absence of position-specific constraint. Additionally, empirically simulated sequence datasets were run through GARD (Kosakovsky Pond et al., 2006), since the method considers both differences in tree topology and branch lengths when calculating the likelihoods of trees. Stark rate heterogeneity could thus easily be mis-interpreted as evidence for recombination by GARD.

### Host-association alleles

In order to test for the presence of alleles associated with host shifts (presumably camel to human) we adapt the 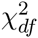 (Hedrick and Thomson, 1986) statistic of LD to estimate the association between host (camel or human) and alleles at polymorphic loci. Briefly, we consider the host to act as a polymorphic site (encoded as H or C, for human and camel, respectively) and compare the association between the “allele” or host and alleles at polymorphic sites. A perfect association of 1.0 could mean, for example, that a biallelic site has one allele that is only found in camel viruses and the other allele only in human viruses.

## Results

### MERS-CoV genome shows evidence of alternative tree topologies

GARD identified a breakpoint at nucleotide position 23722 (corrected ΔAIC=103.6 between single versus two tree model), roughly in the middle of the coding sequence for the S (spike) protein. The 2 phylogenies recovered from this breakpoint were incongruent (see figure S1). Running the resulting fragment 1 (positions 1-23722) and fragment 2 (positions 23723-30126) through GARD again yielded a further breakpoint in fragment 1 at position 12257 (corrected ΔAIC=33.7), near the boundary between ORF1a and ORF1b genes. No more breakpoints could be identified by GARD in the resulting fragments 1.1 (positions 1-12257), 1.2 (positions 12258-23722) and 2 (positions 23723-30126).

5 out of 6 empirical simulation alignments under a nucleotide model simulated without recombination, were identified by GARD as having breakpoints around position 20000, where the clock rate for the rest of the “genome” was increased to be 1.3, 2 or 3 times higher than the first 20kb. Similarly, 2/6 alignments empirically simulated under a codon model were identified as having a breakpoint. Corrected ΔAIC values decreased with decreasing rate heterogeneity, indicating loss of statistical power to detect differences between genomic regions. Analyses in BEAST where the MERS-CoV genome is partitioned into positions 1-23722 and positions 23723-30126 (corresponding to the first GARD-inferred breakpoint) with each partition having an independent molecular clock rate but the same tree or both independent molecular clock rates and independent trees, showed that rate heterogeneity as expressed by the ratio of second fragment rate to first fragment rate to be on the order of 1.513 (95% highest posterior density 1.275, 1.769) for unlinked clocks and 1.375 (95% HPDs 1.079, 1.707) for unlinked clocks and trees (see figure S2). As such, empirically simulated sequence data under 2-fold and 3-fold rate heterogeneity should be considered as caricature examples of a MERS-like organism, which we use to test the sensitivity of the methods we employ.

### MERS-CoV genome exhibits linkage disequilibrium decay

Permutation tests as implemented in LDhat work under the assumption that loci are interchangeable only when there is free recombination or no recombination at all. The tests compare 4 statistics estimated from the actual data to 1000 permutations of the data where site numbers for each locus are reshuffled. Correlation coefficient between two measures of LD, *r*^2^ (Hill and Robertson, 1968) and *D'* (Lewontin, 1964), are expected to show a negative correlation with increasing distance between loci if there is recombination. Permutation of recombining loci will produce a distribution skewed towards more positive values for these two LD statistics and the percentile of the actual observed value can then be used to assess significance.

G4 is the sum of distances between pairs of loci with 4 observed haplotypes, which can only occur if there is repeat mutation or recombination at one of the loci. Under recombination the observed G4 statistic should take a statistically higher value in a distribution of G4 values derived from permuted data. Lkmax is the composite likelihood of pairs of loci under an estimated recombination rate and a given level of sequence diversity. Like the G4 statistic, this statistic is expected to fall in the upper tail of the distribution derived from permuted data in the presence of recombination.

All 4 permutation tests show a consistent signal of recombination in the MERS-CoV genome (see figures 2 and S3). Data from fastsimcoal2 simulations, which did not have rate heterogeneity, produced values for these statistics which mostly fell inside the range of values generated by permuting the simulated data, as expected (figure 2). On 1 occassion this is not the case – simulation 9 passed the Lkmax test and failed the other three. Empirically simulated data, on the other hand, tended to exhibit extreme values, that is the observed value fell below the 2.5th or above the 97.5th percentile of the permuted data, but in ways which were not consistent with recombination. For example, replicate 1 of simulation with 3-fold rate heterogeneity under a nucleotide substitution model exhibits extreme values for all 4 tests, but only one of these – corr(r^2^, d) is consistent with recombination.

### Composite likelihood methods are susceptible to rate heterogeneity

The composite likelihood method, which finds the composite likelihood surface of recombination rate, inferred non-zero recombination rates for all simulated datasets (see figure S4), revealing some degree of susceptibility to both temporal sampling and rate heterogeneity. A window-based approach of this test shows a sharp increase in the recombination rate estimated within 300 nucleotide windows around nucleotide 21000, close to the breakpoint inferred by GARD (see figure S5). We recovered a qualitatively similar pattern when analyzing empirically simulated sequences. It is important to note, however, that none of the simulated data, even under extreme heterogeneity, reproduced the same scale of the estimated recombination rate. Whereas in MERS-CoV data the vast majority of 300 nucleotide windows after position 23000 have a recombination rate per base consistently higher than 0.005, only data simulated under extreme rate heterogeneity approach values as high as that.

In addition to the apparently higher recombination rate in regions with higher rates we expect rate heterogeneity to produce a higher density of polymorphic sites in regions that are evolving faster. This is quite obvious in empirically simulated data with 3-fold rate heterogeneity – the region with higher rate also contains, on average, more polymorphic loci per window in the last third of the “genome” than the first 20kb (see figure S6). We only see hints of this in the actual MERS-CoV genome, with an apparent decline in polymorphism density from position 5000 to 15000 which resembles that of the simulated data with 1.3-fold rate heterogeneity.

### Homoplasies in MERS-CoV genomes are ubiquitous

Homoplasy analyses suggests that the MERS-CoV genome is rife with apparent homoplasies. Both maximum likelihood and Bayesian approaches to ancestral sequence reconstruction converge on similar patterns of homoplasy density (figures 3 and S7). Both methods identify the region around the S (spike) protein as having a high density of synonymous homoplasies.

Empirically simulated sequences showed that homoplasies are not that unlikely in the absence of recombination. All sequences empirically simulated in πBUSS under a nucleotide substitution model had 2-fold homoplasies ranging in frequency from 0.0222 to 0.0550 of all polymorphic sites, with sequences simulated under higher levels of rate heterogeneity having more homoplasies and higher homoplasy degrees (figure 4). However, even under a caricature model of rate heterogeneity we did not reach the same degree of homoplasy as that observed in MERS-CoV, where homoplasic sites comprise as much as 0.1447 of all polymorphic sites and reach homoplasy degrees as high as 4. We were able to recover similar proportions of homoplasic sites from empirical simulations under a codon model (ranging from 0.0821 to 0.1518 of all polymorphic sites), but only under unrealistic values of rate heterogeneity. Even then, the distributions of homoplasy degrees for all simulated datasets are heavily skewed towards low homoplasy degrees, whereas the homoplasy degree distribution for MERS-CoV has a longer tail, indicating that homoplasies are more repeatable across the phylogeny.

Additional tests for excessive homoplasies, PHI and 3Seq also identify MERS-CoV sequences as being recombinant, albeit both spuriously identify some of the simulated datasets as recombinant (see figure S10). Similarly to LDhat results, however, there were no cases where both methods falsely inferred the presence of recombination for the same dataset.

### Model testing supports a model including rate heterogeneity, but not alternative tree topologies

A model including rate heterogeneity alone across breakpoints inferred by the GARD method (*i.e.* linked trees, unlinked relaxed clocks) performs best when applied to MERS-CoV data (figure 5 log marginal likelihoods: -48137.86 and -48138.91, using path and stepping stone sampling, respectively). The next best-performing model (log Bayes factor ≈18) is linked trees and relaxed clocks. Overall, unlinking molecular clock rates between the two genomic partitions appears sufficient to dramatically improve model fit. Additionally, relaxed molecular clocks are preferred over strict molecular clocks (log BF>15, figure 5).

## Discussion

### Recombination tests consistently point to recombination in MERS-CoV

The majority of methods we used (with the exception of marginal likelihood model testing) point to the combined effects of recombination and rate heterogeneity in the genome of MERS coronavirus. GARD (figure 1) identified 2 breakpoints in the genome with high support. It is important to note that calculating the likelihood of a phylogenetic tree involves the estimation of multiple parameters, which at the very least must include a particular tree topology and a set of branch lengths. As such, we considered that the inference of a breakpoint in the MERS-CoV genome could be caused by systematic rate heterogeneity, which we address by evaluating rate heterogeneity across the first breakpoint in BEAST. We estimate an empirical rate heterogeneity ratio between MERS-CoV genome positions 23723-30126 and 1-23722 to be on the order between 1.3 and 1.5 (see figure S2). However, the support for this first breakpoint in MERS-CoV is comparable to support for empirically simulated sequences with 2-fold rate heterogeneity, and breakpoints under MERS-like levels of rate heterogeneity are difficult to detect. Overall, this suggests that evidence for differences in likelihoods between MERS fragments 1 and 2 are beyond what would be expected from rate heterogeneity alone.

**Figure 1.**
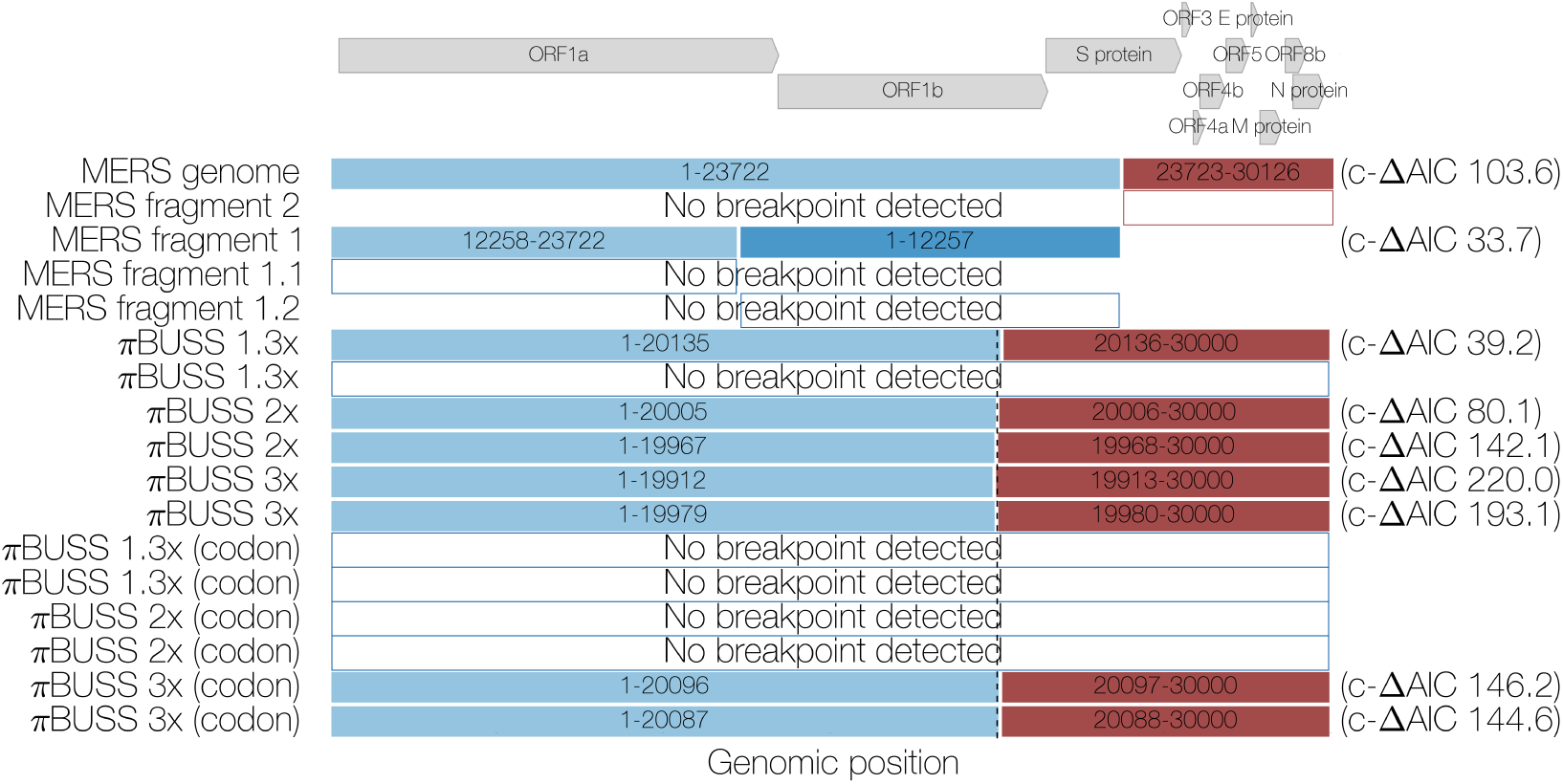
Summary of GARD results. Coloured boxes indicate fragments resulting from GARD-inferred breakpoints with corrected Δ-AIC values shown on the right. Dashed line indicates the actual position where the evolutionary model for simulated sequences under 3 levels of rate heterogeneity is changed. Arrows at the top indicate the positions and names of coding sequences within the MERS-CoV genome.

**Figure 2.**
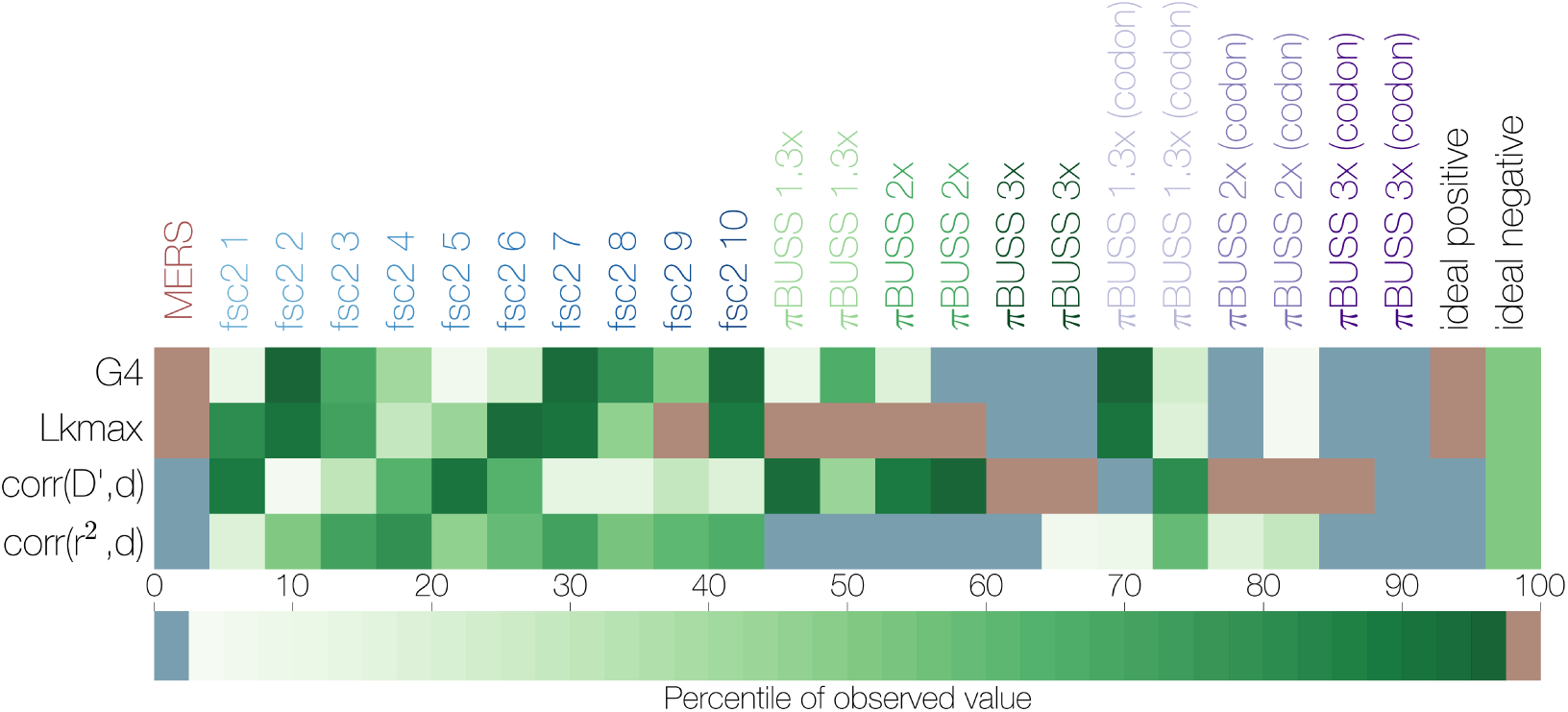
Summary of non-parametric tests for recombination. The percentile of the observed value for 4 statistics of LD decay (y axis) in the distribution of permuted datasets is indicated by colour. Sequence datasets are shown on the x axis, starting with MERS-CoV sequences, followed by 10 fastsimcoal2-simulated datasets and 12 empirically simulated datasets with different degrees of rate heterogeneity. Expected values for ideal datasets are shown in the last two columns, an ideal positive corresponds to the presence of recombination. Values falling between the 2.5th and 97.5th percentile are shown in green, values falling below the 2.5th percentile are in blue, those that are above the 97.5th percentile in red.

Permutation tests show that statistics related to LD decay derived from MERS-CoV sequence data are outliers compared to permuted data (figures 2 and S3). Sequences simulated empirically with varying levels of rate heterogeneity, under nucleotide or codon models of substitution, also have a tendency to exhibit extreme values for these statistics. However, only MERS-CoV data has values for all 4 tests that are in the direction consistent with recombination. For simulated datasets, especially those simulated under extreme levels of rate heterogeneity, values deviating significantly from the permuted data were recovered, but often indicated internally contradictory scenarios.

Homoplasy analyses regardless of inference method show that MERS-CoV sequences contain a large number of homoplasic sites with high homoplasy degrees (figures 3 and S7). Through sequence simulation we also confirmed that both the numbers of homoplasic sites and their homoplasy degrees in MERS-CoV genomes are excessive, even when compared to unrealistic scenarios (*e.g.* 3-fold rate heterogeneity, see figure 4). Homoplasies become increasingly more prevalent when a more realistic codon model is used, due to differences in codon position constraint. Even then, MERS-CoV genomes possess mutations with much higher homoplasy degrees, surpassing simulated datasets with caricature levels of rate heterogeneity. This makes sense under a recombination scenario, as alleles persist in a diverse population and get recombined into novel backgrounds repeatedly, giving an appearance of highly repeatable mutations. Nevertheless, substitution patterns in real genomes are often highly complex and homoplasy-based methods have been shown to be susceptible to rate heterogeneity across sites, especially under higher levels of sequence divergence (Posada and Crandall, 2001). Although rate heterogeneity certainly exists in MERS-CoV data, the divergence levels are still quite low (*θ*/site = 0.0047), giving us some degree of certainty in our inference of homoplasies. It is also reassuring that both maximum likelihood and Bayesian sequence reconstruction converged on similar patterns of homoplasy and synapomorphy across the genome (figures 3 and S7). This is important, since homoplasies inferred using BEAST are integrated over all possible tree topologies, whereas homoplasies inferred by maximum likelihood were conditioned on a single tree. The convergence between these two methods suggests that the data contain enough phylogenetic signal to recover what could be called a “true” tree and that homoplasies, for the most part, can be correctly identified as such.

**Figure 3.**
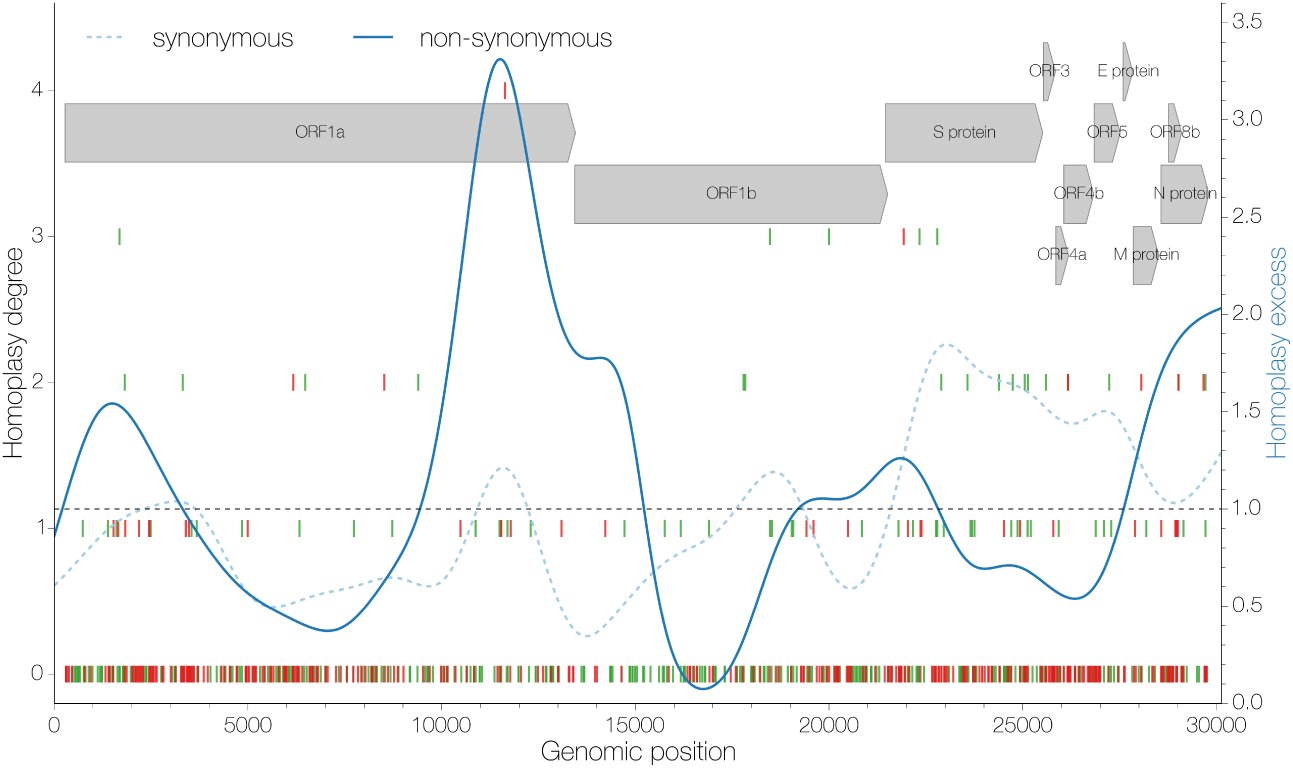
Distribution of apparent homoplasies. Position along the genome is shown on the x axis and homoplasy degree, the number of times a particular mutation has occured in excess in the tree as inferred by maximum likelihood, is shown on the y axis (left). Individual mutations are marked by vertical lines, synonymous ones in green and non-synonymous in red. The ratio of apparent homoplasy over synapomorphy kernel density estimates (bandwidth=0.1) is shown in blue for synonymous (dashed) and non-synonymous (solid) sites separately. Arrows at the top indicate the positions and names of coding sequences within the MERS-CoV genome.

**Figure 4.**
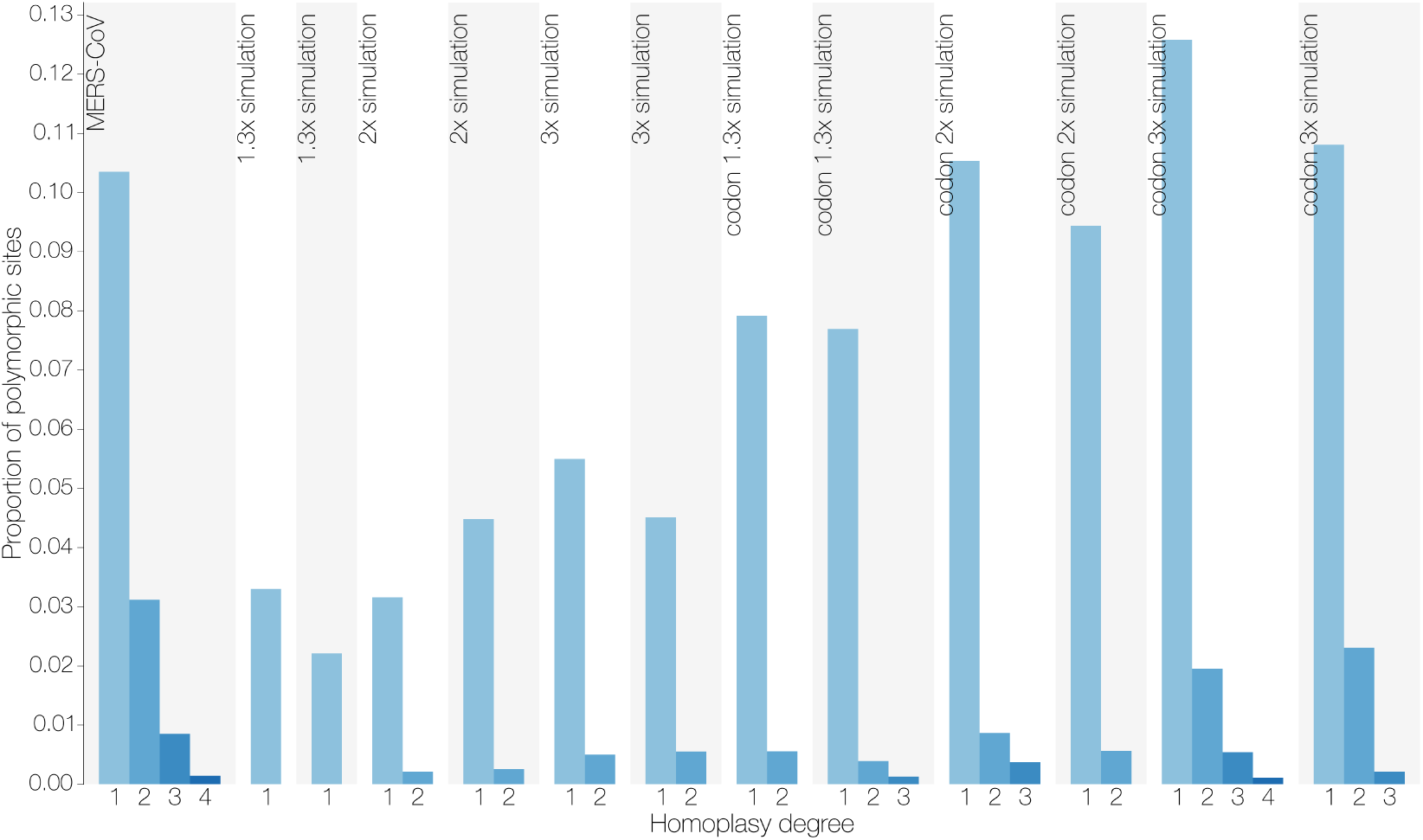
Homoplasy prevalence in MERS-CoV and simulated datasets. Bars show the proportion of all polymorphic sites that are homoplasic, split by homoplasy degree as inferred by maximum likelihood, in MERS-CoV and datasets simulated with different degrees of rate heterogeneity in the presence or absence of site-specific constraint in the form of a codon model. Homoplasy degree indicates how many times a given mutation has occured in excess in the phylogenetic tree.

One major concern surrounding the inference of homoplasies is host adaptation. There are a number of canonical mutations associated with host shifts, *e.g.* the glutamic acid to lysine amino acid substitution at position 627 in the PB2 protein of avian influenza A viruses confers the ability of the virus to replicate in mammals (Subbarao et al., 1993) and a small number of amino acid substitutions in Parvoviruses are associated with adaptations to different hosts (Chang et al., 1992). If MERS-CoV is repeatedly emerging in humans convergent mutations would be expected to arise that might allow the virus to adapt to humans.

However, we expect most host-adaptation mutations to be non-synonymous, whereas we detect both non-synonymous and synonymous homoplasies. This implies the action of recombination, rather than repeated selection for the same host-specific mutations. Furthermore, we do not detect any strong associations between host and particular alleles (figure S8), although we do not believe that there is a sufficient number of sequences from camels to have much confidence in this result.

The overall phylogenetic and genomic patterns of homoplasies are consistent with fairly frequent recombination through time (figure 6). Recent recombination should result in long homoplasy tracts shared across branches in the phylogenetic tree. At most we observe 2 stretches of adjacent homoplasies, one encompassing 3 homoplasies and another encompassing 2 homoplasies, that are shared between taxa, and likely to be caused by recent recombination. The vast majority of homoplasies that we observe, however, occur on their own. Recombination tracts, rather than single template switches are not uncommon in other coronaviruses (Keck et al., 1988; Kottier et al., 1995; Herrewegh et al., 1998). Thus in MERS-CoV we interpret extremely short homoplasy tracts as evidence of relatively frequent recombination. Alternatively, recombination tracts might be short and thus unable to transfer multiple informative sites across lineages.

Unlike all other tests we performed model testing through marginal likelihoods indicates that models including rate heterogeneity explain MERS-CoV data partitioned across a well-supported breakpoint better than models including independent trees. At first this may seem paradoxical, but we believe this result is due to the combined effects of the way homoplasious sites are distributed across the genome and phylogenetic tree of MERS-CoV (figure 6) and the number of parameters involved. A speckled pattern of homoplasious sites without phylogenetic signal could easily be overwhelmed by the signal coming from the sites that support what could be called “the one true tree”, *i.e.* the clonal frame, in the data. Secondly, each phylogenetic tree contains at least n-1 free parameters, so it is not surprising then that models attempting to recover 2 independent trees for both genomic fragments resulting from alternative tree topology analysis of MERS-CoV with highly correlated genealogies are penalized for the extra parameters introduced by a second tree. We would additionally like to point out that the fit of models including relaxed molecular clocks result in dramatic improvements to model fit compared to models with strict molecular clocks (log BF>15, figure 5). Although this could be interpreted as evidence for a considerable degree of lineage rate heterogeneity, the more parsimonious explanation is the ability of a relaxed molecular clock to accommodate homoplasies of recombinant origin, which do not necessarily accumulate at a relatively constant rate like genuine *de novo* mutations do.

**Figure 5.**
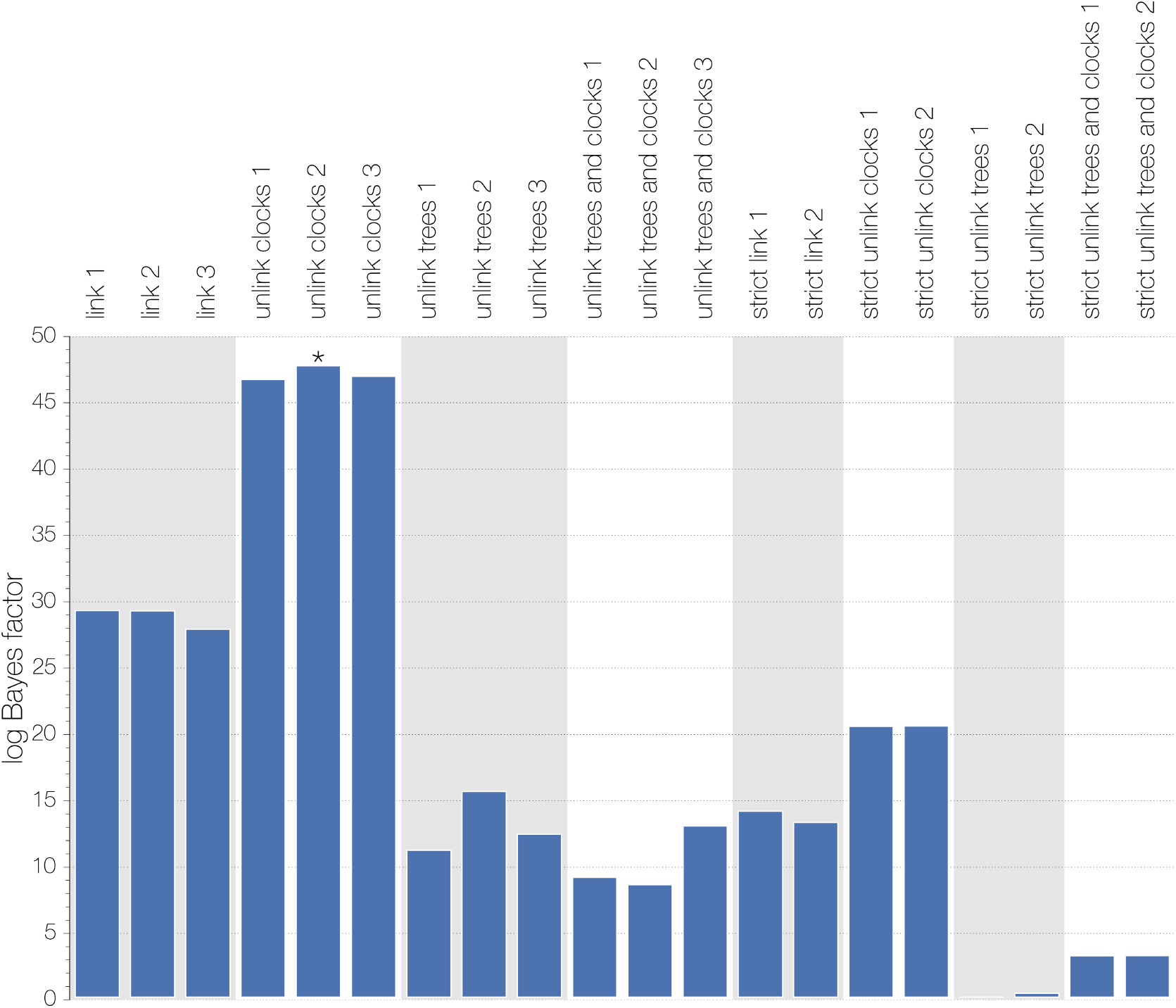
Summary of model comparisons. Difference in marginal likelihoods (Bayes factor) estimated by path-sampling between the worst model (linked strict molecular clock, unlinked trees) and all others. Asterisks indicate the best-performing model (unlinked relaxed clocks, linked trees, run 2) for MERS-CoV data. Analyses employing a relaxed molecular clock were run independently 3 times, those with a strict molecular clock 2 times. Marginal likelihoods estimated using stepping stone sampling gave identical results.

**Figure 6.**
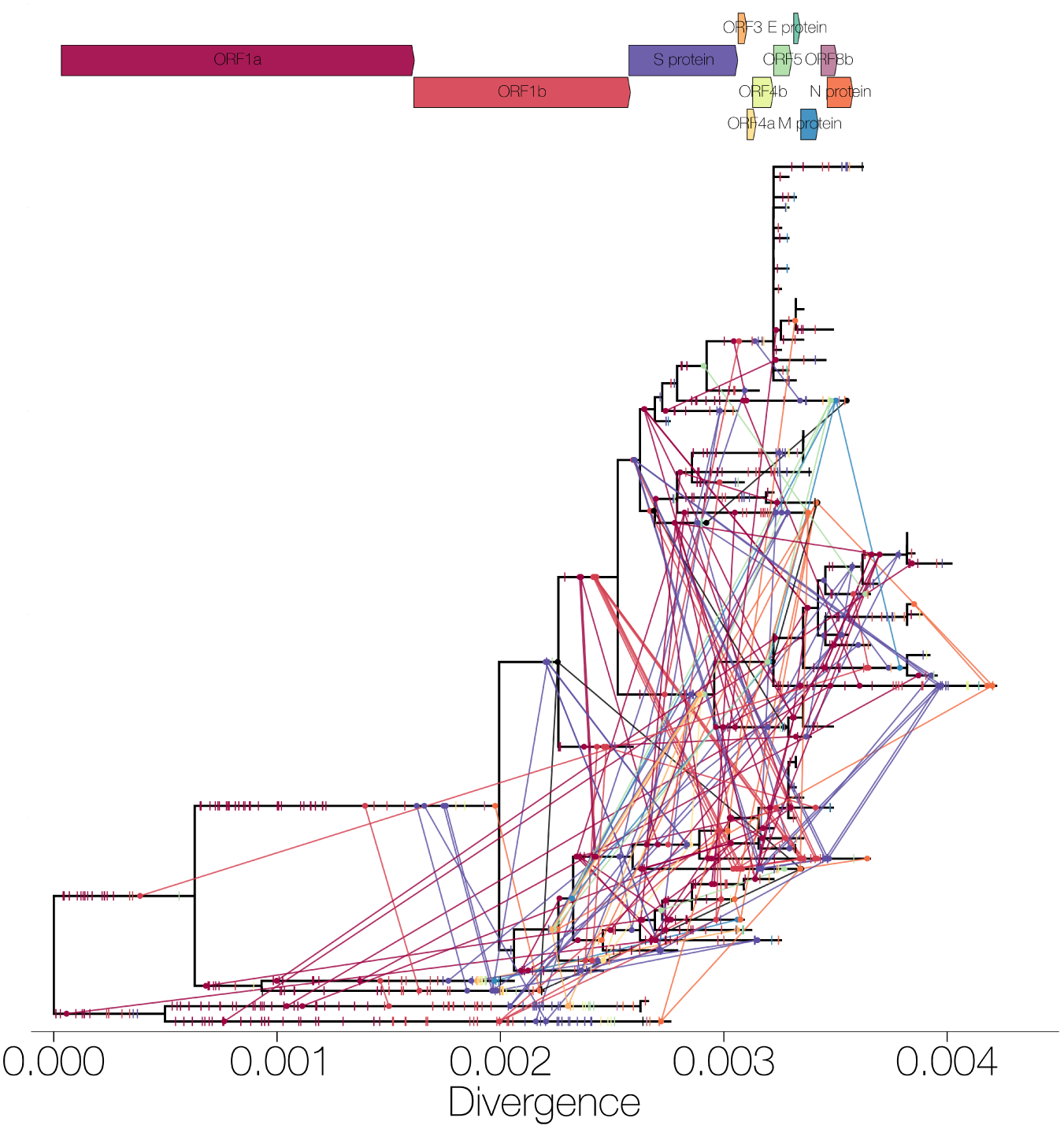
Mutations mapped onto a ML phylogeny. A maximum likelihood phylogeny of 85 MERS-CoV sequences with maximum likelihood-mapped mutations. Synapomorphies are shown as coloured ticks (coloured by coding sequence in which they occur) on branches where they occur. Homoplasies are shown as circles connected with coloured lines, colour corresponds with the coding sequence in which the mutation has occured. Mutations are positioned on the branches in proportion to where the mutation occurs in the genome, *e.g.* mutations shown towards the end of a branch correspond to mutations near the 3’ terminus of the genome. Arrows at the top indicate the order, relative length and names of coding sequences within the MERS-CoV genome.

### Implications for future analyses

Recombination aside, MERS-CoV genomes exhibit a significant degree of rate heterogeneity amongst sites. Marginal likelihood analyses indicate that estimating independent molecular clocks after partitioning the MERS-CoV genome into 2 fragments alone substantially increases model fit over a completely linked (trees and clocks) model (log Bayes factor ≈18). This highlights the advantage of employing relaxed molecular clocks, as in our case the method is clearly capable of accomodating recombination in an otherwise entirely clonal analysis framework. In addition, previous studies of SARS-like coronaviruses in bats have identified recombination breakpoints in small numbers of isolates falling close to the “transition zone” around site 22000 (Hon et al., 2008; Lau et al., 2010) which in our analysis of MERS-CoV is where GARD, LDhat and BEAST identify changes in the underlying model of evolution (figures 1, S5 and 3). Overall, a more detailed investigation will need to be done to determine if empirical patterns of rate variation in MERS-CoV have the potential to generate apparent recombination signals.

We also expect that as more sequences of MERS-CoV become available more homoplasies will be detected, some contributing to the homoplasy degree of the homoplasies already reported here, some previously unknown and some turning mutations currently thought of as synapomorphies into homoplasies. Although new sequences are likely to come from human cases, we think that sequencing MERS-CoV circulating in dromedary camels is of extreme importance from both surveillance and epidemiological points of view.

### Implications about the virus population structure and infection dynamics

Our results point towards frequent recombination in MERS-CoV in the recent history of the MERS-CoV outbreak. For this to occur different lineages of the virus must encounter each other often and implies frequent co-infection with MERS-CoV. To date it is difficult to ascertain whether the human infections with MERS-CoV are a result of substantial asymptomatic transmission amongst humans, or repeated zoonosis of the virus from camels to humans or a combination thereof. Given the severity of MERS we find it unlikely that humans could be sufficiently frequently co-infected with two or more different lineages of the virus. Previous serological studies have failed to find evidence of prevalent past MERS-CoV infections of humans (Gierer et al., 2013; Aburizaiza et al., 2013), although a recent nation-wide study in Saudi Arabia has detected non-negligible numbers of individuals with antibodies against MERS-CoV, especially amongst shepherds and slaughterhouse workers (Müller et al., 2015). We thus propose that MERS-CoV mostly infects, and recombines, in camels. A study by Adney et al. (2014) has shown that camels only suffer mild symptoms from MERS-CoV infection and numerous other studies indicate an extremely high prevalence of antibodies specific against MERS-CoV in camels (Müller et al., 2014; Corman et al., 2014b; Chu et al., 2014; Reusken et al., 2013, 2014). At the same time, however, sequencing has not indicated the presence of multiple infection in camels, or any other animal. We believe that individual MERS-CoV co-infections are rare, but given the size of the epidemic in camels, as inferred from serology, the total number of co-infections is high. In addition, MERS-CoV infection is transient in camels Adney et al. (2014) and thus sequencing efforts, which have been insufficient and very limited in camels, are highly unlikely to capture a co-infection.

Another point worth considering is that alleles that have arisen through mutation in MERS-CoV can be recombined, increasing the genetic variation of the virus (Muller, 1932). Whether this is of epidemiological importance for humans depends entirely on what alleles are circulating in the reservoir, although there is no evidence that MERS-CoV is particularly likely to become as transmissible as common human pathogens or even SARS-CoV.

### Data availability

Python scripts used to process trees and sequences are available at:

https://github.com/evogytis/MERS_recombination/tree/master/scripts.

Input and output files for programs used are publicly available at:

https://github.com/evogytis/MERS_recombination.

## Acknowledgements

GD was supported by a Natural Environment Research Council studentship D76739X. The research leading to these results has received funding from the European Research Council under the European Community’s Seventh Framework Programme (FP7/2007-2013) under Grant Agreement no. 278433-PREDEMICS and ERC Grant agreement no. 260864.

